# Host age at experimental *Helicobacter pylori* infection shapes epithelial development of mouse gastric organoids

**DOI:** 10.1101/2025.08.31.672357

**Authors:** Hjördís Birna Árnadóttir, Danai Anastasia Panou, Jorge F. L. Teixeira, Felix Romer, Katrine B. Graversen, Sigri Kløve, Sandra B. Andersen

## Abstract

The bacterium *Helicobacter pylori* colonises gastric glands, which triggers direct responses to bacterial activity and genetic modifications of the host. While most frequently asymptomatic, infection can cause stomach cancer through a stepwise sequence from chronic gastritis to carcinoma, involving induction of stem cell-like properties of epithelial cells. We tested the effect of host age at infection on epithelial development in a mouse organoid model, where mice were infected as neonates or adults, for one month. To isolate the effects of host genetic modification, gastric organoids were grown in the absence of *H. pylori* and sequenced and imaged. We found that *H. pylori* infection early in life resulted in a larger size of the derived organoids. The expression of marker genes in organoids for different cell types was dependent on host age, suggesting a decrease in pit cells and an increase in endocrine cells with age. *H. pylori* infection early in life accelerated this age dependent shift, and we propose that the cell type profile affects the host response to infection.

## Introduction

Stomach infection with *Helicobacter pylori* induces a chronic inflammatory response that, in a subset of individuals, progresses to peptic ulcers or gastric cancer. While most infections result in mild, asymptomatic gastritis, approximately 10-15% of individuals experience ulcers, and 1% develop gastric cancer ^1^. This makes *H. pylori* the primary cause of an estimated 90% of cases of gastric cancer worldwide, which remains the fifth leading cause of cancer-related mortality ^2^. Malignant transformation typically follows Correa’s cascade, a stepwise sequence beginning with chronic active gastritis and advancing to dysplasia and ultimately to carcinoma ^3^. Hallmarks of this transition are the loss of gastric glands, replacement of native epithelial cells by intestinal-like cells, and induction of stem cell-like properties in differentiated epithelial cells ^3,4^. The host effects arise from a combination of direct responses to bacterial activity, and genetic modifications of the host. These may be challenging to disentangle when analysing patient samples or samples from experimental *in vivo* infection.

The carcinogenic potential of *H. pylori* is attributed to DNA double-stranded breaks, which can result in chromosomal mutations by faulty repair, and epigenetic modifications such as promoter hypermethylation that silences tumour suppressor or DNA repair genes ^5^. *H. pylori* inflicted damage may thus leave a lasting imprint on the host genome, affecting the tissue even after the infection is eradicated with antibiotics ^5^. However, not all infected individuals follow this pathogenic trajectory, and disease outcome is shaped by a complex interplay between host genetics, bacterial traits, and the match between host and strain genetics ^6,7^. The timing of *H. pylori* infection is another factor ^8–10^. Early-life acquisition is expected to be the norm in high prevalence populations, and experimental early-life infections in mice tend to induce immune tolerance, which reduces the risk of T-cell driven auto-immune diseases. Here the same virulence factors contributing to disease, may confer protective effects ^11^. Late-life infections often provoke a more robust immune response, resulting in greater inflammation and associated disease risks ^10^. The effect of age at infection on the host epithelial response has, however, not been studied in detail.

Work with gastric organoids is instrumental for understanding *H. pylori* - host interactions at the cellular level ^4,12–14^. Gastric organoids and derived monolayers have been used to show that *in situ* expression patterns can be replicated *in vitro*, maintaining epigenetic changes ^15,16^. We use mouse gastric organoids derived from adult tissue-residing stem cells to investigate how the timing of *H. pylori* infection influences the imprint on the host epithelial response, in the absence of bacteria. To model early- and late-life infection, organoids were established from mice infected for four weeks, starting either at 11 days or at seven weeks of age and compared to organoids derived from uninfected mice. We hypothesised that infection-derived organoids would exhibit altered morphological features compared to controls, driven by the cell-proliferative effects of *H. pylori* infection. Furthermore, we wanted to investigate whether late-life infection would elicit a stronger pro-inflammatory response compared to early-life infection. We found that early life infection had a cell proliferative effect, resulting in larger organoids. Further, we observe that the antrum cell type composition is age dependent, and that infection induces lasting changes to gastric cell biology that shifts the profile towards that of older tissue.

## Methods

### H. pylori culturing for animal infection

Wild type PMSS1 *H. pylori* ^*17*^ was grown from frozen stock on TSA Sheep Blood plates (Thermo Fisher Scientific) in an incubator (HERACELL VIOS 150i, Thermo Fisher Scientific) at 37 °C, 6% O_2_ and 10% CO_2_. After 72 h, bacteria were collected and resuspended in 1 ml of Brucella broth (BD Biosciences), and used to inoculate 20 ml of Brucella broth media, containing 10% heat-inactivated foetal bovine serum (FBS, Sigma-Aldrich) and 0.06 mg/ml vancomycin (Sigma-Aldrich), and grown overnight with 100 rpm shaking. Then 10 ml media were added to the cultures, and incubation continued for 2 – 3 h. The motility of the bacteria was confirmed, after which the cultures were centrifuged at 2000 rpm for 10 min at room temperature. A bacterial pellet from 20 ml culture was resuspended in 500 µl culture media. OD_600_ was measured, and serial tenfold dilutions inoculated on blood agar plates for colony counting after 72 h.

### Animal experiments

For the early-life (EL) infection, C57BL/6J mice from Janvier (France) were set up for breeding with two females and one male per cage. Eleven days after birth, pups from two litters were inoculated by oral gavage with ∼ 50 µl of 10^9^ CFU *H. pylori* PMSS1 or sterile Brucella media. Mice were weaned at 3.5 weeks of age and separated by sex. For the late-life (LL) experiments, six week old C57BL/6J mice were purchased from Janvier and inoculated at seven weeks of age by oral gavage with 100 µl bacterial culture. Mice were inoculated on two consecutive days and euthanised 30 – 44 days later for collection of the antral part of the stomach tissue. The first batch of EL organoids (EL1) were grown using tissue from two control mice pooled together (2 female) and two infected mice pooled together (1 male, 1 female; all five weeks old). In EL2 we used tissue from two control mice (not pooled; 1 male, 1 female) and two infected (1 male, 1 female; all seven weeks old). In LL1 and LL2 we used tissue from one control (male) and one infected (male) each (Fig. S1). The experiments were done on different days. Mice were maintained in individually ventilated cages at 22 ± 1 °C on a standard 12 hour light-dark cycle with *ad libitum* access to water and standard chow. Mice were euthanised by cervical dislocation after inhalation of isofluran, and the stomach dissected out, opened along the lesser curvature, and the antral part sectioned. Tissue for gland isolation was processed immediately, while tissue for DNA extraction was stored at −20 °C. Experimental procedures were approved by the Danish Animal Experimentation Inspectorate (2020-15-0201-00432).

### DNA extraction from stomach tissue for H. pylori screening

DNA was extracted from mouse stomach tissue with the DNeasy Blood & Tissue Kit (Qiagen) to assess whether the *H. pylori* inoculation was successful. Bead beating was used to lyse the tissue instead of an overnight incubation step. The presence of *H. pylori* in the stomach samples was checked with PCR, using primers targeting a region upstream of the OipA gene ^18^. Included in the PCR was a positive and a negative control. Bacterial load was quantified with qPCR and normalised to total DNA concentration as previously described ^18^. Unfortunately, no tissue from EL1 control animals were stored for DNA extraction.

### Methylation sequencing

Reduced Representation Bisulphite Sequencing was performed at Novogene UK on the DNA from stomach sections. Data was analysed in nf-core/methylseq pipeline (v3.14.0) ^19^ using bwa-meth align with the methylkit flag to generate files for further analyses in RStudio with methylKit ^20^, genomation ^21^ and GenomicRanges ^22^. Reads were filtered for low count = 10 and high percent = 99.9, normalized by the median, and only CpG with standard deviations larger than 2% were kept.

### Isolation and seeding of murine gastric glands for organoid culture

Isolation and culture of murine gastric glands were performed as previously described with slight modifications ^12, 16^. Briefly, fresh murine antral tissue was kept on ice in basal DMEM/F12 (Gibco) medium supplemented with Glutamax (Gibco), HEPES (Gibco) and Primocin (Invitrogen). After removing the overlying mucus and outer muscle layers using forceps, the tissue was washed in cold chelating buffer (PBS (Gibco), sucrose, D-sorbitol (Sigma-Aldrich/Merck), and DL-dithiothreitol (Sigma-Aldrich/Merck), cut into two – five mm^2^ pieces and washed vigorously. The washed tissue was incubated in chelating buffer supplemented with 10 mM EDTA at room temperature for 10 min. Glands were released by applying pressure in a petri dish, collected in cold basal medium and separated from tissue debris by settling and centrifugation (200 x g, 5 min at 4 °C). Glands were pelleted and kept on ice. For seeding, glands were resuspended in Cultrex type 2 (R&D Systems) on ice and 50 μl was placed in the center of each well of a pre-warmed 24-well plate creating a dome Following a brief incubation, domes were overlaid with 500 µL of gastric organoid medium (basal medium supplemented with 3dGRO L-WRN conditioned media, Sigma-Aldrich), gastrin (Phoenix Pharmaceuticals), EGF and FGF-10 (R&D Systems), N-acetylcysteine (Sigma-Aldrich/Merck), B27 (Invitrogen), and the ROCK inhibitor (Invitrogen). Organoids were cultured at 37 °C in 7% O2 and 5% CO2 (HERACELL VIOS 160i, Thermo Fisher Scientific). RHOKi was removed 2 days post-seeding, and the media was refreshed every two to three days. In two wells for each EL group no organoids were left at day six after seeding and these were excluded from subsequent analyses.

### Optical imaging of gastric organoids

Organoids were imaged on day six after seeding while still intact in the 24-well plates with an Evos XL Core inverted microscope with 4× magnification, resulting in four – six images per well. The focus was adapted so that as many organoids as possible were in focus in any given image. The brightness settings remained unchanged between images. MATLAB (R2023b) was used to analyse optical images of the organoid cultures with the OrganoSeg script ^23^. The same parameters were applied to all images; intensity threshold: 0.453, window size: 100, and size threshold: 340. The organoid area (in pixels) was calculated and then converted to µm^2^, and the median organoid area was calculated for each well and log-transformed. Dead organoids and debris that got segmented by the software were manually picked out from each picture by one individual.

### Whole-mount immunofluorescence staining and imaging of gastric organoids

After six days of culture, organoids were fixed in 4% paraformaldehyde followed by permeabilization with 0.1% PBS-Tween. Organoids were blocked with an organoid washing buffer (OWB) containing Triton X-100 (Thermo Fisher), and BSA (Sigma-Aldrich) in PBS and transferred to 15 ml tubes after centrifugation (200 x g, 4 min, 4 °C). The organoids were stained with Alexa Fluor 647 Phalloidin (1:400 dilution, Invitrogen) and DAPI (1:1000 dilution, Invitrogen), in OWB at room temperature for 1.5 hours, washed twice, and resuspended in Vectashield (Vector Laboratories). 50 μl of the suspension was mounted on slides sealed with MOLYKOTE grease (DuPont) and stored at 4 °C.

Imaging was performed on a CorrSight Spinning Disk confocal microscope (FEI) using a 10× objective. Whole-slide maps were generated with widefield mode, and individual organoids were imaged using Z-stacks acquired with the spinning disk function. The number of planes was calculated based on the diameter of each organoid: the location of the approximate top and bottom was identified and the diameter of the organoid was then divided by the appropriate Nyquist frequency. For imaging we used channels SD405 for DAPI and SD640 for Alexa Fluor 647 Phalloidin. From the EL group, we imaged 75 organoids from uninfected controls and 67 organoids from *H. pylori* infected mice, of these 37 and 23, respectively, were from EL1.

Confocal images were stitched and tiled together using Fiji ImageJ (v2.9.) with a custom Fiji macro ^24^. Huygens Professional (v23.04) was used for deconvolution of images for clearer fluorescent signals. The same parameters were applied to all pictures. Imaris (v10.1) was used for cell counting and diameter measurements. Larger organoids were folded, which reduced image quality in the bottom section. Hence, the log-transformed number of cells in the top half of each organoid were used, counted from the largest diameter. Spot counting parameters were set manually for each organoid due to varying signal intensity. Fiji ImageJ was used for editing figures.

### Optical and spinning disc confocal image data analysis

Linear mixed-effects models (LMMs) were performed with lme4 ^25^ in RStudio (ver. 4.3.2) ^26^ to assess the effects of infection and age at infection on organoid size. Infection treatment and age at infection were included as fixed effects, while biological replicate (animal) was included as a random effect to account for inter-animal variability. Wells were treated as technical replicates. Visually inspecting the residual plots showed no major deviations from normality or homoscedasticity. Post hoc testing with Tukey’s HSD was used to analyse differences between experimental groups in the optical data set with emmeans ^27^. For the optical data, a LMM was used to test if organoid size was correlated with number of organoids in a well by experimental group, with biological replicate as a random effect. The medians of log-transformed cell counts and cell count per area for EL samples were compared with Wilcoxon tests.

### RNA preparation and sequencing from gastric organoids

RNA was extracted from gastric organoids with the RNAqueous – Micro Kit (Invitrogen), following the provided protocol. In short, 100 – 200 μl lysis solution was added directly to each well and the Cultrex dome was scraped into a sterile Eppendorf tube for each technical replicate. The tubes were vortexed and placed at −20 °C until further processing. The samples were thawed and 100% ethanol was added to a v/v of 66.6% and the sample was vortexed. RNA was eluted twice with 10 μl of pre-heated Elution Solution. The Qubit RNA HS Assay Kit (Invitrogen) and Qubit 3 Fluorometer were used to measure RNA concentration. RNA quality was measured with the Bioanalyzer (Agilent) using the Agilent RNA 6000 Nano Kit. Samples were sent for mRNA sequencing at Novogene UK on the Illumina NovaSeq paired-end 150 bp platform.

Transcriptome data was analysed in nf-core/rnaseq pipeline (v3.14.0) ^19^. Trimming and filtering was done using fastp ^28^. Poly-G filtering was done additionally. Removal of technical duplicates was carried out using dedup (highest accuracy score, 6). The nf-core/rnaseq pipeline was used with default settings which uses STAR ^29^ to align reads to the mouse reference genome GRCm39 release no. 109 (https://useast.ensembl.org/) and Salmon ^30^ to quantify mapped reads and generate a gene length scaled count matrix.

Differential gene expression analysis was performed in RStudio using DEseq2 ^31^ on the count matrix scaled by gene length. Genes with reads present in less than four samples were filtered away prior to analysis. The differential gene expression analysis was done in three parts. We looked at the effect of mouse age by contrasting control early life and late life. We then looked at the effect of treatment, where the design input was time of infection and treatment group, contrasting *H. pylori* and control. Next, we combined time and treatment in a four-level factor and contrasted experimental groups, *H. pylori* against control, in EL and LL. Heatmaps of DEGs were made with pheatmap ^32^. We used STRING enrichment analysis (version 12.0; https://string-db.org/; (42)) to assign differentially expressed genes (DEGs) to known functional classes and cellular pathways, using the list of expressed genes as statistical background. Additionally, we used Microsoft Copilot to search for relevant patterns in the gene lists.

### Statistical analyses

In Rstudio we used tidyverse ^33^, ggplot2 ^34^, ggpubr ^35^, gridExtra ^36^, in addition to the abovementioned packages. Statistical significance was defined as *p* < 0.05.

## Results

### H. pylori infection status

Infection status of the mice was confirmed by PCR of DNA isolates from stomach tissue in all but the LL1 *H. pylori* mouse, which was therefore excluded from the analyses. QPCR data indicated that the bacterial load was 9748 *H. pylori* gene copies/µg DNA in the LL2 sample, compared to a median value of 9966 *H. pylori* gene copies/µg DNA in the EL animals (range 2391-17374 *H. pylori* gene copies/µg DNA, where the samples with the outlier values were from EL1 used for the organoids that were RNA sequenced).

### Methylation patterns and exclusion of data from EL2 experiment

We obtained a median ± MAD of 58.2 M ± 5.45 M reads per sample, which after nf-core processing was reduced to 1.65 M ± 78.6 K methylated sites per sample. A PCA plot of the samples confirmed that the LL1 *H. pylori* treated animal, which was found to be uninfected, was most similar to LL1 and LL2 control samples (Fig. 1). Most notably, however, was that the EL2 samples were furthest away from all other samples, and less similar to each other across treatment. Given this, and the lack of EL1 control samples available for DNA extraction, we could not perform statistical analyses on the differences in methylation patterns. In the transcriptome data presented below, results from EL2 were also outliers (see below). We decided to exclude data from this experiment, and therefore also present the microscopy data without EL2. Animals from EL2 were from the same litter as EL1, but euthanised two weeks later. We speculate that conditions at the animal housing facility could have changed in this time, contributing to the differences.

**Figure 1:**
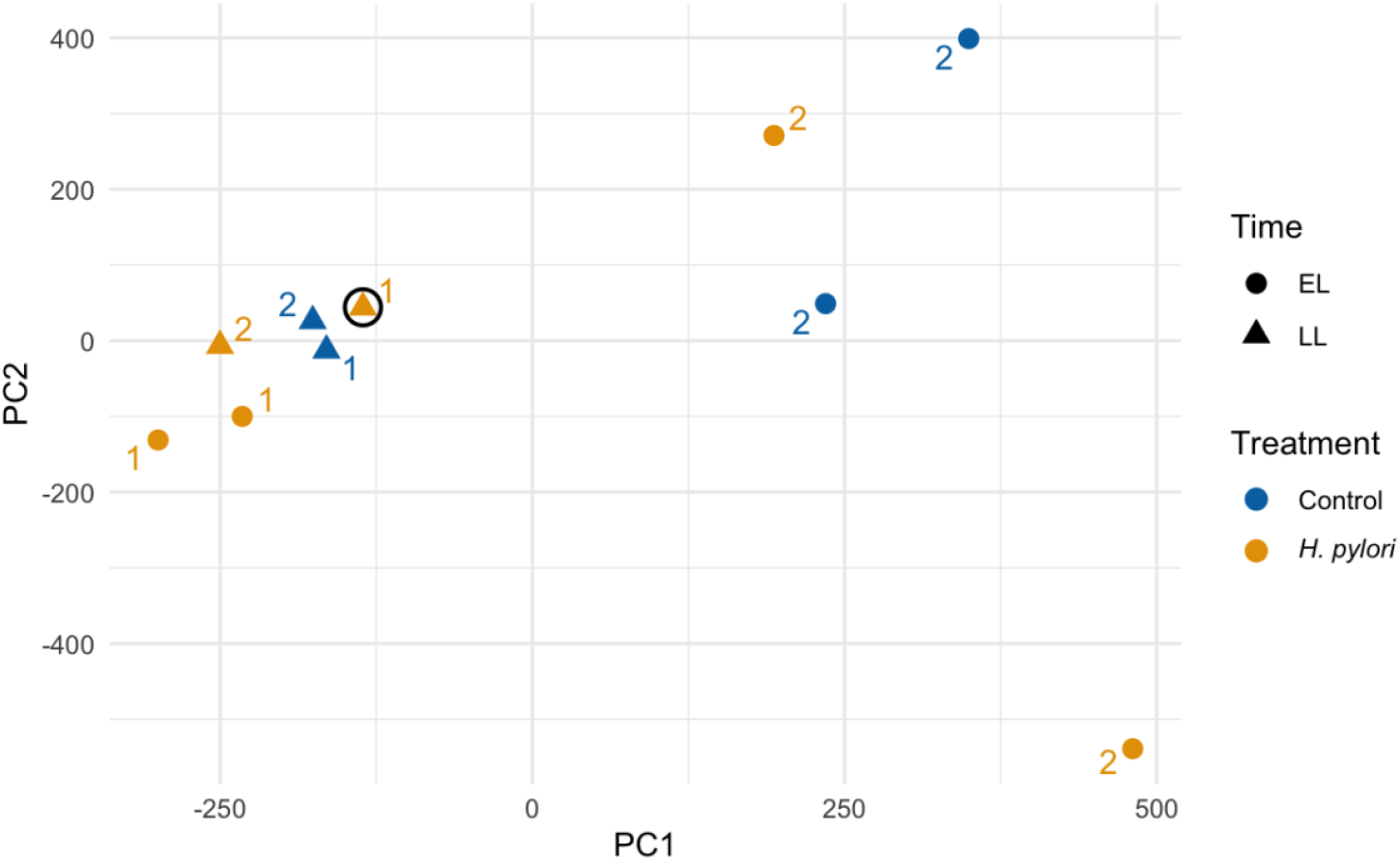
PCA plot of the methylation patterns of EL1, EL2, LL1 and LL2 experiments. Color indicates the treatment (Control or H. pylori), shape the time (EL or LL) and number the experiment (1 or 2). The LLHp1 sample that came from a mouse that turned out to be uninfected is highlighted with a circle, and grouped with the samples from control mice.

### Organoids from mice infected with H. pylori early in life are larger than all others

We found a significant effect of *H. pylori* infection on the log-transformed median organoid size (LMM: β = 1.03, SE = 0.18, t_36_ = 4.06, *p* = 0.0003), and an interaction between infection status and time (LMM: β = −1.23, SE = 0.39, t_36_ = −3.15, *p* = 0.03, overall R^2^ = 0.35), where organoids derived from EL *H. pylori* mice were significantly larger than those from EL control mice (Post hoc test, *p* < 0.0014; Fig. 2). The number of glands was not standardized per well before seeding, but did not vary significantly across groups (Two-Way ANOVA; F_1, 38_ = 1.62, *p* = 0.21), and we did not find a significant association between the number and the size of organoids grown in the same well (LMM: β = −0.02; SE = 0.013; t_36.95_ = −1.59; R^2^ = 0.071; *p* = 0.12). This was therefore not included as a variable in the analysis. Including data from experiment EL2 did not change the outcome of the analysis.

**Figure 2:**
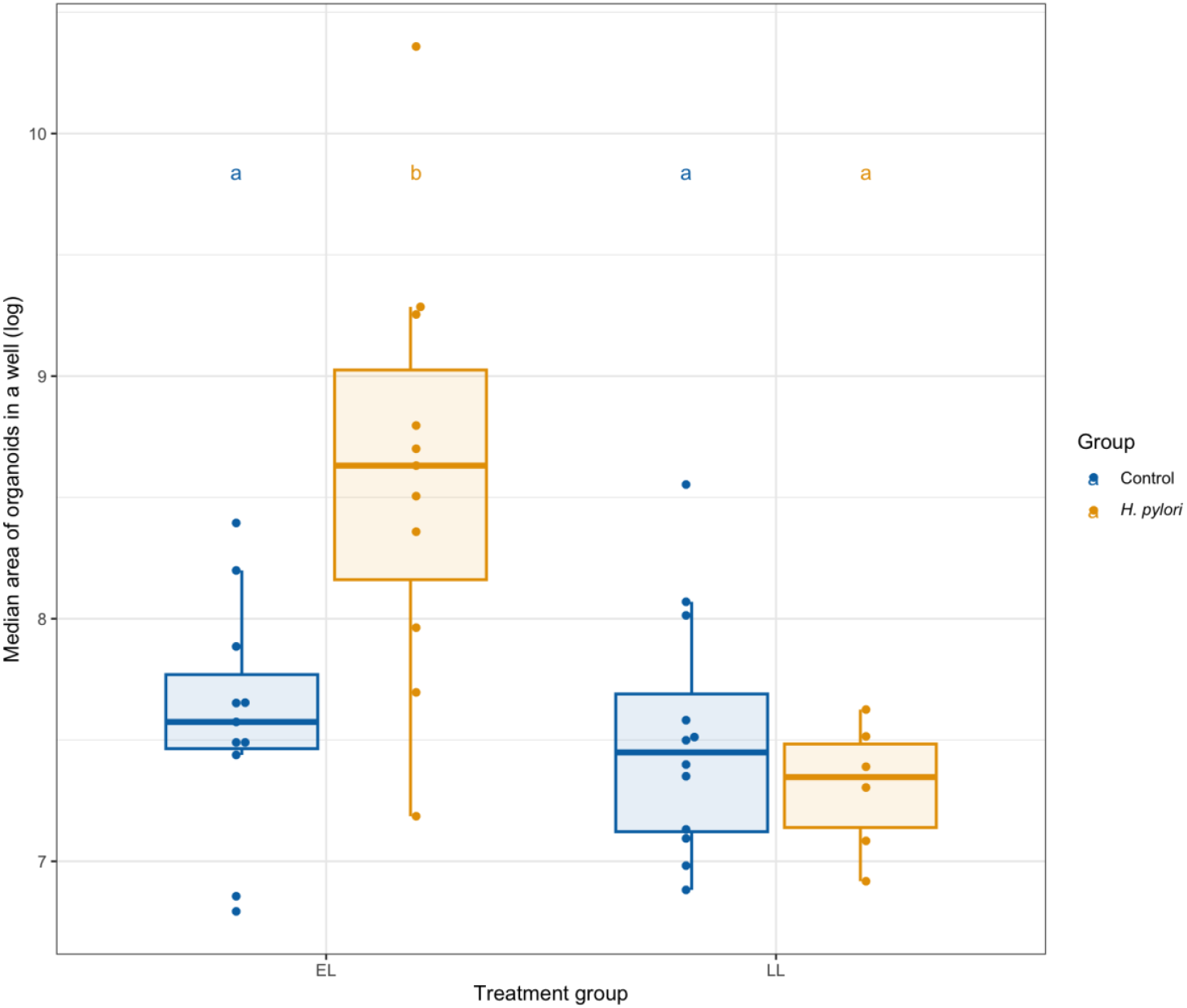
Organoids from mice infected early in life were significantly larger than those from early-life control mice. Each data point on the boxplot represents the median organoid size in each well. Tukey HSD post hoc test results are indicated with letters (a and b), showing that EL H. pylori organoids were significantly larger than EL control organoids, while there was no statistically significant difference between LL organoids.

### Organoids from mice infected with H. pylori early in life contain more cells

We found that the significantly larger organoids from mice infected early in life also contained more cells than those from control animals (2.8× more cells on average, Wilcoxon test on log-transformed cell count number in top-half organoid: W = 224; *p* = 0.0018; Fig. 3). Additionally, there were significantly more cells per area in the organoids from infected mice (1.5× more cells per area on average; Wilcoxon-test on log-transformed cell count per area in top-half organoid: W = 258; *p* = 0.010). Cell numbers varied substantially across samples, ranging from 9 to 6870 cells, reflecting heterogeneity in growth of organoid culture. To validate the use of top-half counts as a proxy for total cell number as the larger organoids were collapsed, we analyzed a subset of organoids that could be fully imaged. A strong linear relationship was observed between top-half and total cell counts (also including data from EL2 to increase sample size; adjusted R^2^ = 0.960; F(1,12) = 316.1, *p* = 5.49×10^−10^), with a slope of 1.73. While this indicates slight overestimation relative to the expected factor of 2, the approach was deemed robust for smaller organoids. However, in larger organoids with deep epithelial folds, top-half estimates may be less accurate due to imaging limitations.

**Figure 3:**
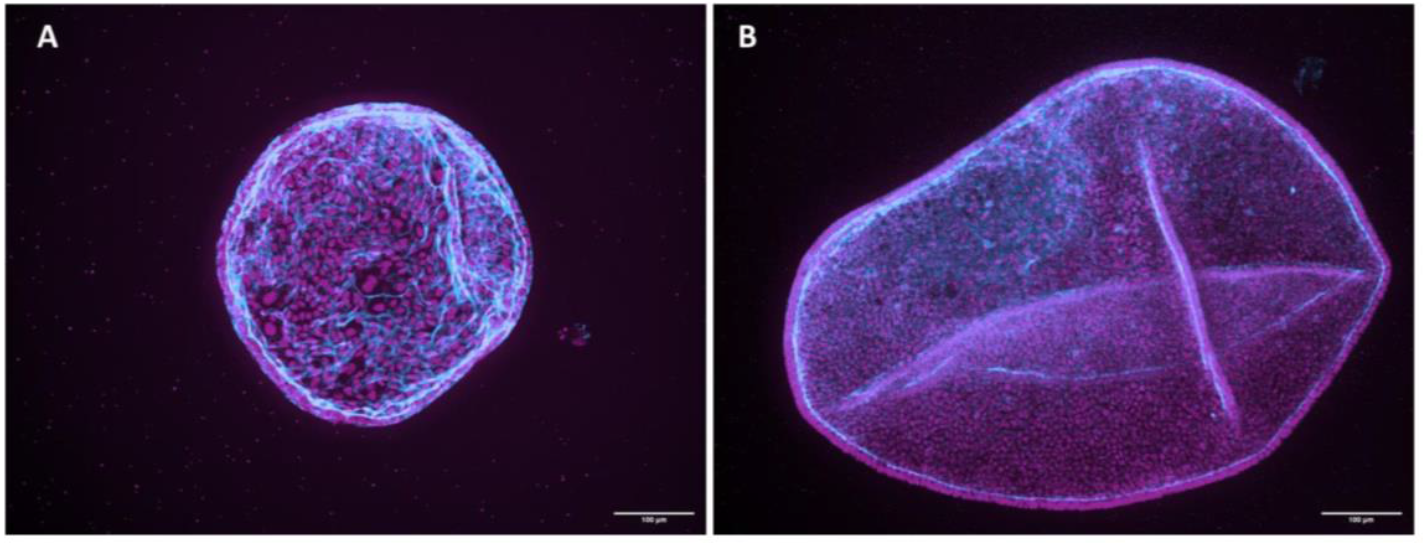
Representative spinning disk confocal images of early life control and infection-derived organoids. A) Early life control organoid, 756 cells in top half. B) Early life infection-derived organoid, 3532 cells in top half. Samples were stained with Alexa Fluor 647–conjugated phalloidin for filamentous actin (F-actin) and DAPI for nuclei. Images acquired using a 10× objective; scale bar: 100 µm.

#### Age-dependent transcriptomic responses to *H. pylori* infection in gastric organoids

Sequencing of mRNA from seven day old organoids provided a dataset with a median ± MAD of 37.3 M ± 3.8 M reads per sample. Following processing in nf-core 15.4 ± 1.7 M reads per sample scaled by gene length were passed to DESeq2 analyses for normalisation and analyses of differential gene expression. From the EL2 experiment, data was only obtained from one control and five *H. pylori* technical replicates. Initial PCA analysis demonstrated that the two early life experiments, i.e. the biological replicates, exhibited greater transcriptomic divergence compared to late life experiment replicates, which clustered tightly (Fig. 4A). Because the EL2 control sample lacked technical replicates it was not possible to include Experiment as a factor in the DESeq2 analyses. As the EL2 experiment was also an outlier in the methylation data (Fig. 1), data from this experiment was excluded from the subsequent analyses, with cautioning of the interpretation of the results.

**Figure 4:**
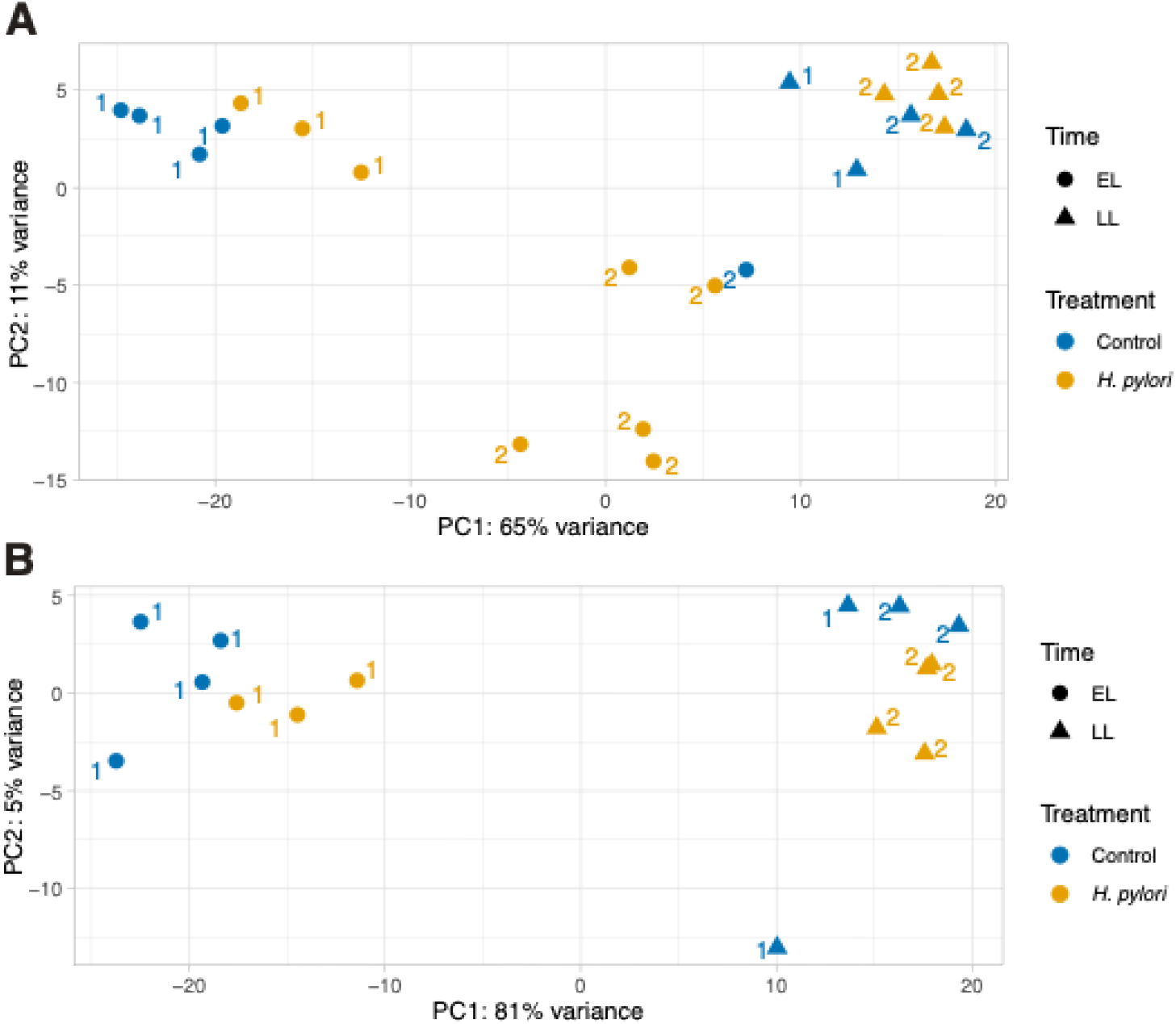
Principal component analysis (PCA) for RNA sequencing data. A) PCA plot for all biological replicates included in RNA sequencing. B) PCA plot after EL2 was removed from the analysis.

Filtering decreased the number of genes included in the analysis from 57,010 to 22,116. Excluding the outlier biological replicate improved the separation between conditions in the PCA, with PC1 and PC2 accounting 81% and 5% of the variance, respectively (Fig. 4B). As also indicated by the PCA plot (Fig. 4B), transcriptomic profiling revealed that the most pronounced gene expression differences occurred between early and late life organoids, where 2484 genes had lower and 2845 genes higher expression in early life compared to late life control samples.

We identified 222 unique differentially expressed genes (DEGs, Table S1-S4), of which 201 exhibited normalised read counts >50 in at least one sample and were included in a heatmap (Fig. 5, Fig. S2). In the analyses we also included the gene *Ghrl* that encodes the gastric hormone ghrelin with *p*_adj_ = 0.085, as expression is known to be affected by *H. pylori* infection. By hierarchical clustering we grouped these DEGs into seven expression clusters. Notably, EL control organoids displayed a transcriptionally distinct profile, whereas *H. pylori* infected organoids - regardless of infection age - clustered more closely with late-life samples. This indicates an infection-associated convergence in gene expression towards resembling samples from the LL category. Of these, 120 genes had the highest expression in EL control (cluster 3 and 4), and 25 genes had the lowest expression (cluster 1). STRING enrichment analysis of these clusters revealed significant over-representation of genes involved in biological processes such as biotic stimulus and lipid metabolism, and KEGG pathways related to drug metabolism (all genes except one also covered in the biotic stimulus process), NOD-like signalling (all genes also covered in the biotic stimulus process), and fat digestion and absorption (all genes also covered in lipid metabolism processes, Fig. 5). In contrast, genes in cluster 2, 5, 6 and 7 were differentially expressed in *H. pylori* infected organoids, either across both ages or in an age-specific manner or only in early or late life infections. In these, no processes or pathways were found to be enriched by STRING analyses.

**Figure 5:**
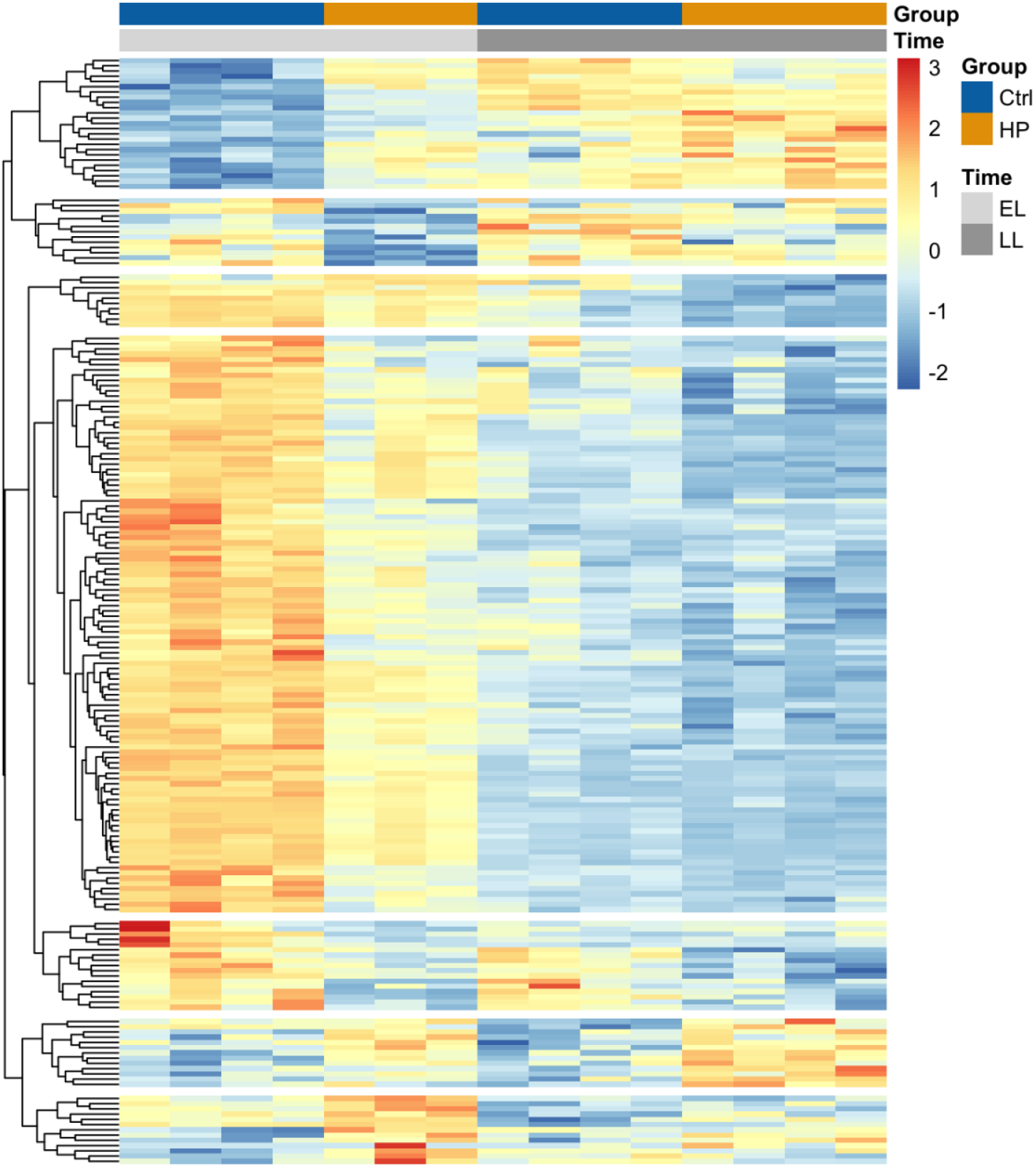
Heatmap of scaled expression of genes that were differentially expressed with a p_adj_ < 0.05 in the analyses; contrasting control and H. pylori organoids, EL control and H. pylori organoids, and LL control and H. pylori organoids. Hierarchical clustering was set to seven which resulted in well-resolved clusters with similar expression patterns. Heatmap is annotated with gene names in Fig. S2.

#### Expression of markers for cell types differ over time and infection

DEGs related to stomach epithelial development and mucus composition suggest that the cell type population changed with age and infection. Expression of the genes *Muc5ac, Gkn1, Gkn2*, and *Tff1* and *Psca*, which are markers of mature and immature pit cells, respectively ^13,37,38^, decreased with a median of 90% with age (comparing control EL to control LL), and was a median of 53% lower in infection derived organoids (comparing EL control to EL *H. pylori*, and LL control to LL *H. pylori*; Fig. S3). Expression of the transcription factor genes *Klf4* and *Grhl3*, that affect expression of the pit cell markers, and genes for collagen secreted by the pit cells, *Col4a1* and *Col4a2* as did the enzyme gene *Gsta1* and *Mucl3* marker of immature pit cells (Fig. S3). Additionally, so did the *Lgr4* gene that has been shown to have the highest expression in isthmus stem cells ^39^, although not significantly so. Expression of *Ces2a, b* and *c*, which have been found to be expressed from chief cells, also declined with age, and infection. This was in contrast to the genes for the gastric hormones gastrin, PYY and ghrelin, the protein chromogranin A and the histamine decarboxylase enzyme, which are secreted by enteroendocrine cells ^38^. Overall, these had a median of 643% increased expression with age that was altered in infection derived organoids (150% increased for ghrelin, and 34% decreased for gastrin and PYY), while EL infection caused an increased expression of *Chga* (276%) and *Hdc* (250%) and a decrease in LL infection (25 and 28 %, compared to controls, Fig. S4). Expression of the base stem cell markers *Lgr5, Axin2, Sox9* and *Aqp5* increased median of 250% with age but was not affected by infection status, while *Muc6* and *Tff2* expression was consistent across time and treatment ^39–41^.

#### Developmental and inflammatory responses to *H. pylori* infection

Several genes that have been found to link differential growth, inflammation and cancer exhibited interesting expression patterns. A cluster of genes centered on the transcription factors *Irf7* and *Stat1* follows the expression pattern of pit cells markers and could thus be expressed by these, with the highest expression in EL, decreasing with *H. pylori* infection and age (Fig. S4). These are involved in type I interferon-based responses to host-microbe interactions. Crosstalk from *Irf7* and *Stat1* can affect expression of *Nfkbia*, which is a negative regulator of the NF-κB complex that can dampen inflammation ^42^, and follows the same pattern (Fig. S5). In contrast, expression of the *Ikbkg* gene, which is a central hub activating *Irf7* and NF-κB ^43^, was upregulated in *H. pylori* organoids independent of age (Fig. S6).

Also independent of infection time, the genes *Lama5* and *Bcl2*, which are indirectly linked in the STAT3 pathway that promote gastric cancer development ^44,45^, were upregulated. In contrast, expression of *Syk*, which is positively correlated with survival of gastric cancer patients ^46^, was downregulated in infection. Expression of the growth factor *Fgf9*, that has been implicated in promoting invasibility in gastric cancer ^47^ and can be involved in activation of Wnt signalling ^48^, and the Wnt ligand *Wnt5a* was relatively low but increased by *H. pylori* infection (Fig. S6).

Organoids derived from early life infection, which were significantly larger than the others, showed differential expression of genes related to cell death and differentiation (*Xkr8, Vmp1* and *Rpl35a*), developmental signalling (*Epc1*), and inflammation and metabolism (*Hpgd* and *Pla2g1b*), however, none of these were obviously connected in pathways to other differentially expressed genes (Fig. S7).

A small subset of genes were differentially expressed only in late life infection-derived organoids. Of them, *Pxdn, Tmem176a, Rab15* and *H2*.*Ab1* were upregulated in LL *H. pylori* organoids, and *Atf3* was downregulated. The peroxidase encoded by *Pxdn* contributes to extracellular matrix stability by crosslinking collagen, and is involved in inflammatory responses ^49^. Increased expression is a biomarker of gastric and other types of cancer ^50^. Methylation of the transmembrane protein gene *Tmem176a* has been linked to lymph node metastasis of gastric cancer, where expression is negatively correlated with patient survival rates ^51^. Expression of the transcription factor *Atf3* is stress-induced, and while it can both promote and suppress inflammation and apoptosis ^52^, it is considered to be tumor suppressing ^53^. Its reduced expression has been observed in cancerous tissues compared to healthy ^53^. Expression of the antibacterial protein *Reg3g* decreased with age but was maintained in LL *H. pylori* organoids. As it targets gram-positive bacteria it may contribute to favouring *H. pylori* in the stomach ^54^ (Fig. S8).

## Discussion

While host age at infection is known to affect the outcome of experimental *H. pylori* infection, the age-dependent impact of *H. pylori* infection on gastric epithelial growth and transcriptional profiles has not been studied in detail. The strongest effect of *H. pylori* was observed following EL infection. The organoids from EL infected mice were significantly larger than organoids from all other groups (Fig. 2 and 3). These large organoids contained a significantly higher number of epithelial cells, and more cells per area, suggesting that EL infection exerts a pronounced and lasting influence on gastric epithelial cell morphology and architecture. While such structural alterations are consistent with the current literature that *H. pylori* disrupts cell-cell junctions and reorganizes the cytoskeleton ^55,56^, there were only a few genes where expression in EL *H. pylori* organoids was significantly different from that of the other groups. Rather, EL *H. pylori* organoids displayed transcriptional profiles for the DEGs primarily resembling those of the LL organoids, irrespective of host infection status (Fig. 5).

Mouse age at sacrifice had the largest effect on the global gene expression profile, and differential expression of markers for specific cell types suggests a shift in epithelial composition. To our knowledge such a change with age has not been documented before. We observed that EL gastric organoids exhibited higher levels of pit cell markers (*Muc5ac, Gkn1, Gkn2*, and *Tff1* and *Psca*). The higher expression of pit cell markers in EL organoids is in line with data from adult human samples, showing that pit cells were rare in 3D organoids under the same growth conditions as we employed, without withdrawal of Wnt-3α ^13,57^. In contrast, there was an enrichment of markers for basal enteroendocrine and stem cells in LL organoids. An increase in *Lgr5* expression with age has also been observed in rats ^58^, but we found no effect of infection despite *H. pylori* previously being shown to induce hyperproliferation of *Lgr5*+ stem cells ^4,14^. We speculate that infection causes an acceleration of tissue development, with a shift in cell types. This is in line with our recent study comparing gene expression of stomach tissue from EL infected and control mice. Here we found that one week after infection the tissue expression profile of *H. pylori* infected mice already resembled that of samples collected after two weeks, when looking at genes related to the extracellular matrix, muscle contraction and metabolism, ^59^.

*H. pylori* shows a tropism for pit cells ^13^, thus young mice having more of these cells could contribute to the larger effect of EL infection, as observed for size and transcriptome. The basal gland cells, for which marker genes were more abundantly expressed in late life samples, initiate the inflammatory response to *H. pylori* infection, and could thus contribute to the more extensive immune reaction that has been observed to late life infection ^60^. The infectious burden of the host did not play an obvious role in our study, even though other studies have shown it to be higher for mice infected early in life, compared to late in life, one month after infection ^60,61^. We found that the *H. pylori* burden of the late life infected mouse to be intermediate to that of the four mice infected early in life.

The inflammatory host response to *H. pylori* infection contributes to cell proliferation ^3^. Several of the genes that were differentially expressed in organoids from infected animals, have been found to link inflammation and cancer development. For the genes associated with interferon response and NF-κB signalling, the infection response appeared to be related to the hypothesised shift in cell types.

As the organoids themselves were not infected with *H. pylori* it implies that any observed effects are either epigenetic or caused by host mutations, acquired during the initial host infection. This has previously been observed for *H. pylori* infection, with sequencing of infected tissues and in organoids derived from patients ^15^. Unfortunately, our biological sample size was too small to explore this in detail, but in a PCA plot of methylation patterns the samples segregated based on age and infection status (Fig. 1).

### Limitations and challenges

A limitation of the study is the small number of biological replicates, and the exclusion of data from EL2. In addition, both male and female mice were used for the EL experiments but only male for the late life experiments. While not explored in detail for *H. pylori*, there may be sex differences in immune response to the infection. RNA was extracted from all organoids in a well, regardless of their size and condition. The larger size of early life infection-derived organoids may affect the transcriptional profile, as organoids of different sizes could be at different stages of development and cell differentiation. This may potentially be addressed by sequencing organoid-derived 2D cultures, although the profile between 2D and 3D cultures can also differ ^13^. We did not have enough biological replicates to identify specific methylation differences, however, Fig. 1 suggests that epigenetics are likely to contribute to the age and infection derived differences we see in gene expression. Several of the genes we found to be differentially expressed have previously been shown to be affected by epigenetic changes in other *H. pylori* studies and given the short experimental time such changes are perhaps more likely to occur than host mutations.

## Concluding remarks

The model used in this study is well suited for investigating the effects of infection on host cells in terms of their morphology and gene expression. Length of infection could also be implemented into the model, for instance to study further changes in bacterial load or gene expression. By using this model, it is possible to isolate the effect of infection on the gastric tissue, in the absence of bacteria and the immune system. Additionally, tissue cultures could be re-exposed to *H. pylori* to explore differences in response. As organoid or organoid-derived monolayer infection studies can only address effects of acute infection, using tissue from infected animals may be a way to incorporate long-term effects of infection. The results also clearly indicate that host age at sacrifice matters for the cell composition of the resulting cell culture.

## Data availability

Transcriptome and methylation data will be available as NCBI BioProject ID PRJNA1311342.

## Author contributions

HBA & JFLT performed the experiments, HBA, KGB, DPA, FR, JFLT, & SBA performed the data analyses, JFLT & SBA designed the experiments, HBA, DPA & SBA drafted the manuscript, and all authors contributed to revisions.

## Supplementary material

**Figure S1:** Schematic figure showing the experimental work flow. Created in BioRender. Andersen, S. (2025) https://BioRender.com/mxw2nr1

**Figure S2:** Heatmap of scaled expression of genes that were differentially expressed with a p_adj_ < 0.05 in the analyses; contrasting control and *H. pylori* organoids, EL control and *H. pylori* organoids, and LL control and *H. pylori* organoids. Hierarchical clustering was set to seven which resulted in well-resolved clusters with similar expression patterns.

**Figure S3:** Organoid expression of genes that are markers of pit cells, with read counts normalised to gene length on the y-axis. Boxplots depict the median and the interquartile range, whiskers the min- and max values with individual data points as dots. Note the different scales on the y-axis. The x-axis shows the infection age as early-life (EL) and late-life (LL) and color denotes the treatment as either control (blue) or *H. pylori* (yellow).

**Figure S4:** Organoid expression of genes that are markers of endocrine cells, with read counts normalised to gene length on the y-axis. Boxplots depict the median and the interquartile range, whiskers the min- and max values with individual data points as dots. Note the different scales on the y-axis. The x-axis shows the infection age as early-life (EL) and late-life (LL) and color denotes the treatment as either control (blue) or *H. pylori* (yellow).

**Figure S5:** Organoid expression of genes linked to *Irf7* and *Stat1* interferon signalling, that follow the expression pattern of the pit cell markers. Read counts are normalised to gene length on the y-axis. Boxplots depict the median and the interquartile range, whiskers the min- and max values with individual data points as dots. Note the different scales on the y-axis. The x-axis shows the infection age as early-life (EL) and late-life (LL) and color denotes the treatment as either control (blue) or *H. pylori* (yellow).

**Figure S6:** Expression of genes linked to cell proliferation that are differentially expressed in *H. pylori* organoids, with read counts normalised to gene length on the y-axis. Boxplots depict the median and the interquartile range, whiskers the min- and max values with individual data points as dots. Note the different scales on the y-axis. The x-axis shows the infection age as early-life (EL) and late-life (LL) and color denotes the treatment as either control (blue) or *H. pylori* (yellow).

**Figure S7:** Organoid expression of genes that are differentially expressed in early life *H. pylori* organoids, with read counts normalised to gene length on the y-axis. Boxplots depict the median and the interquartile range, whiskers the min- and max values with individual data points as dots. Note the different scales on the y-axis. The x-axis shows the infection age as early-life (EL) and late-life (LL) and color denotes the treatment as either control (blue) or *H. pylori* (yellow).

**Figure S8:** Organoid expression of genes that are differentially expressed in late life *H. pylori* organoids, with read counts normalised to gene length on the y-axis. Boxplots depict the median and the interquartile range, whiskers the min- and max values with individual data points as dots. Note the different scales on the y-axis. The x-axis shows the infection age as early-life (EL) and late-life (LL) and color denotes the treatment as either control (blue) or *H. pylori* (yellow).

**Table S1:** Differentially expressed genes comparing early-life control organoids to late-life control organoids. Output from DESeq2, listing statistics testing how gene expression changes with age, by contrasting early-life control organoids and late-life control organoids. Genes with an adjusted *p*-value < 0.05 were considered to be statistically significantly differentially expressed.

**Table S2:** Differentially expressed genes comparing infection-derived organoids to controls (Hp-effect analysis). Output from DESeq2, listing statistics testing how gene expression changes with infection, by contrasting control organoids and *H. pylori* organoids. Genes with an adjusted *p*-value < 0.05 were considered to be statistically significantly differentially expressed.

**Table S3:** Differentially expressed genes comparing early-life control organoids to early-life *H. pylori* organoids (Early life analysis). Output from DESeq2, listing statistics testing how gene expression changes with infection early in life, by contrasting early-life control organoids and early-life *H. pylori* organoids. Genes with an adjusted *p*-value < 0.05 were considered to be statistically significantly differentially expressed.

**Table S4:** Differentially expressed genes comparing late-life control organoids to late-life *H. pylori* organoids (Late life analysis). Output from DESeq2, listing statistics testing how gene expression changes with infection late in life, by contrasting late-life control organoids and late-life *H. pylori* organoids. Genes with an adjusted *p*-value < 0.05 were considered to be statistically significantly differentially expressed.

