## Supplementary figures and images for "Host age at experimental *Helicobacter pylori* infection shapes epithelial development of mouse gastric organoids"

### Fig.s1

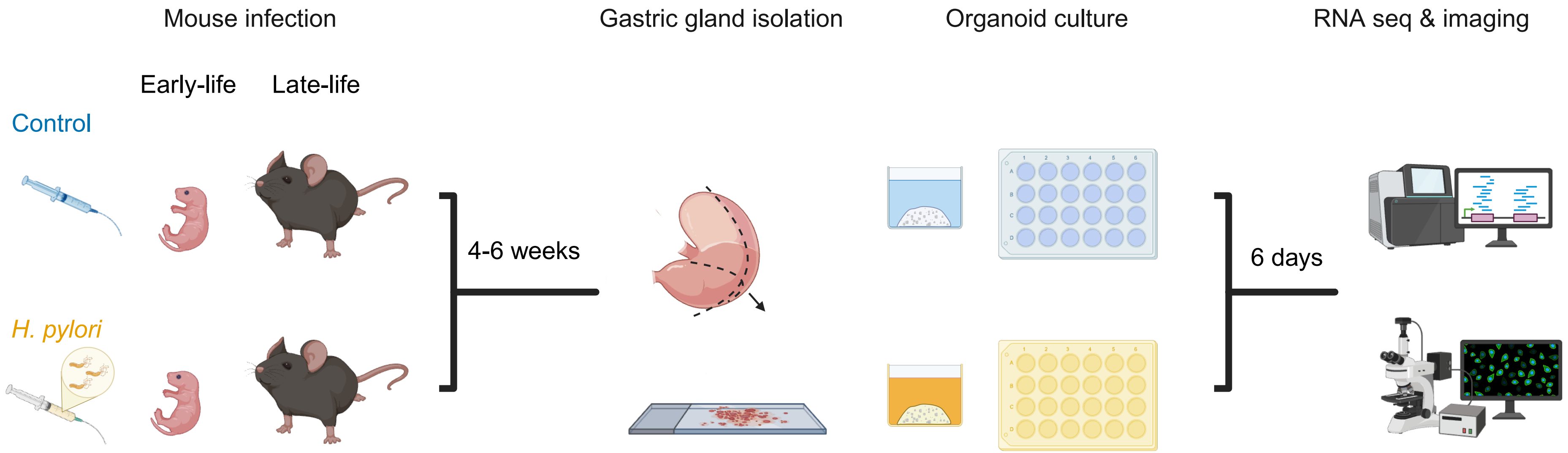

### Fig.s2

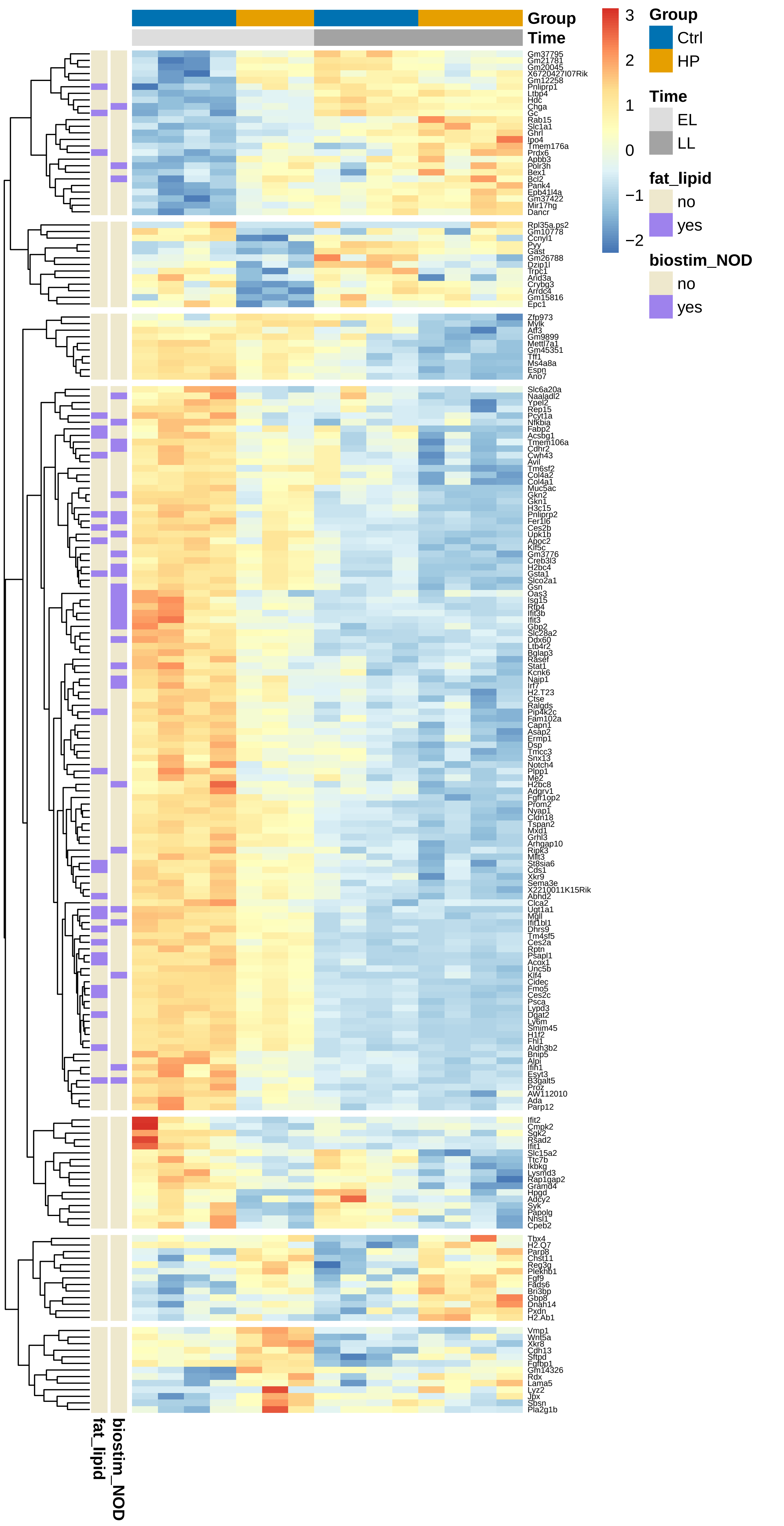

### Fig.s3

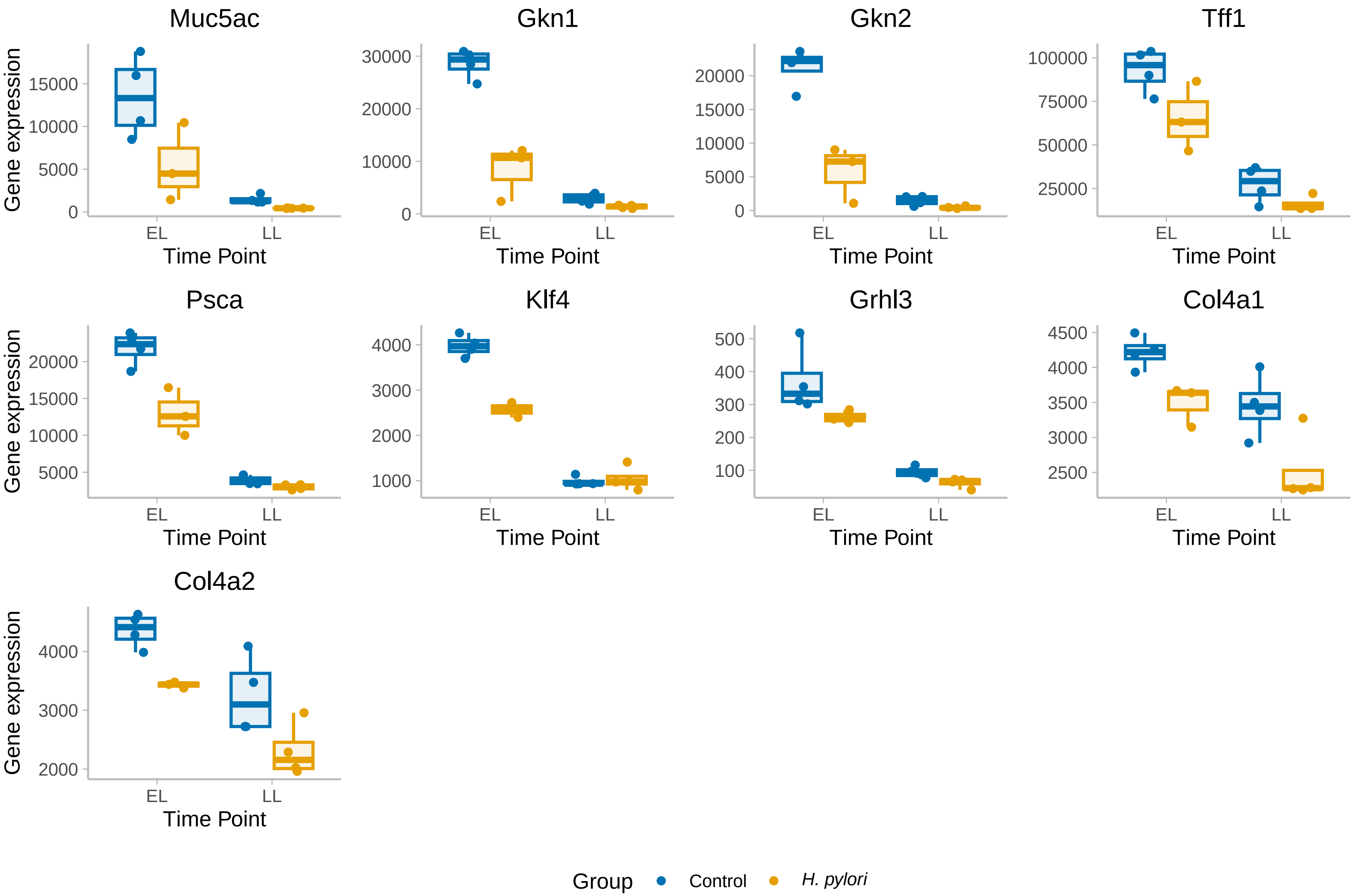

### Fig.s4

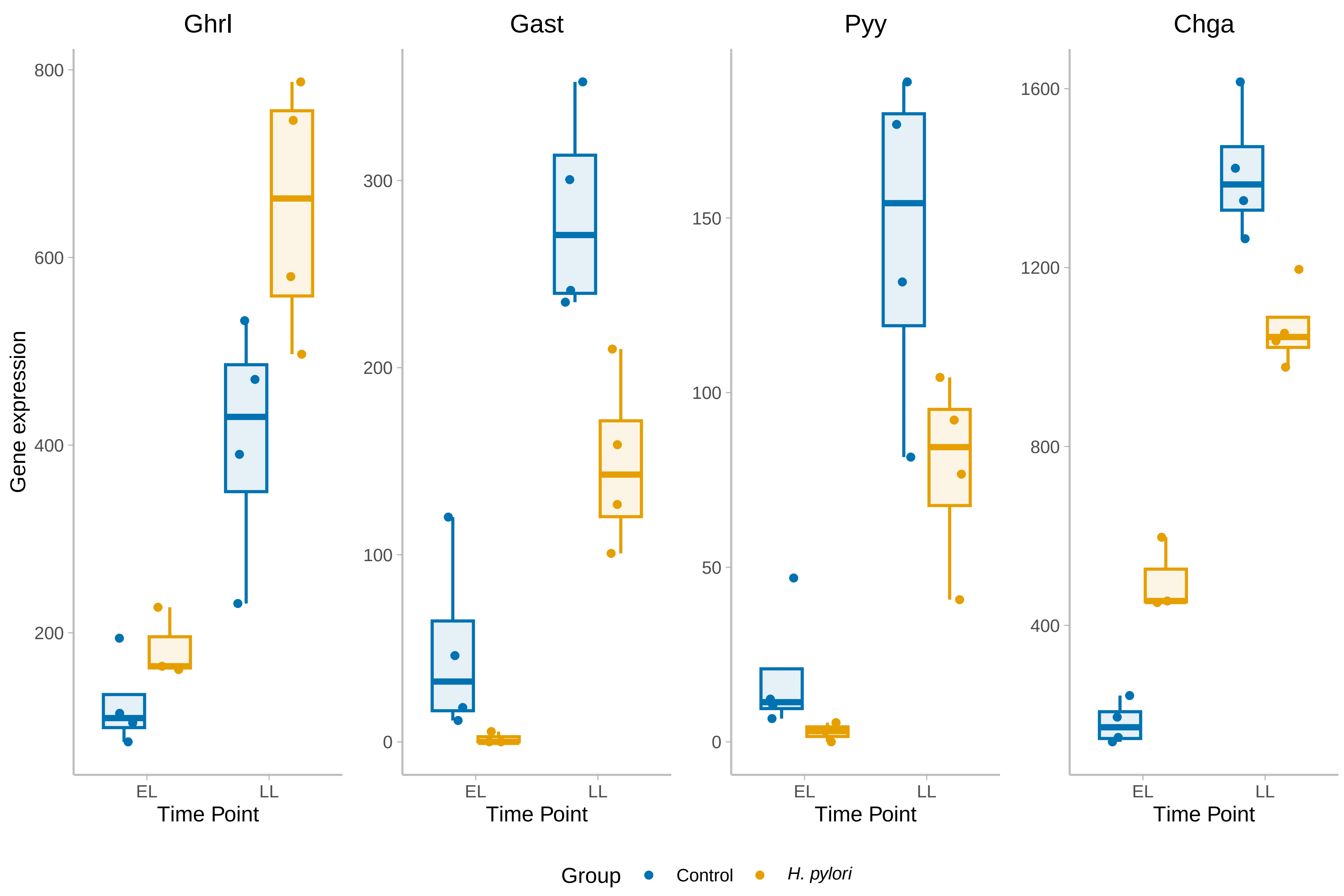

### Fig.s5

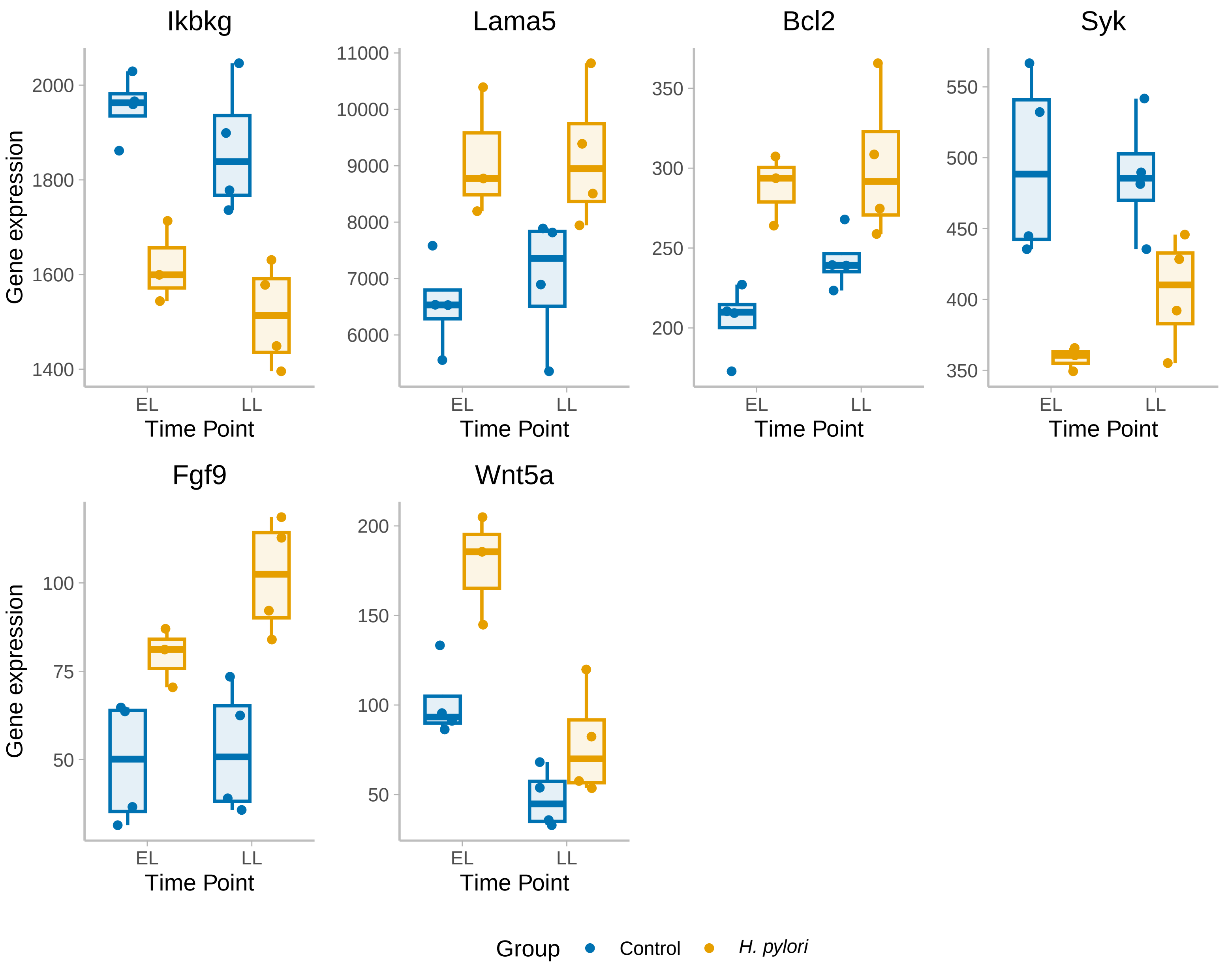

### Fig.s6

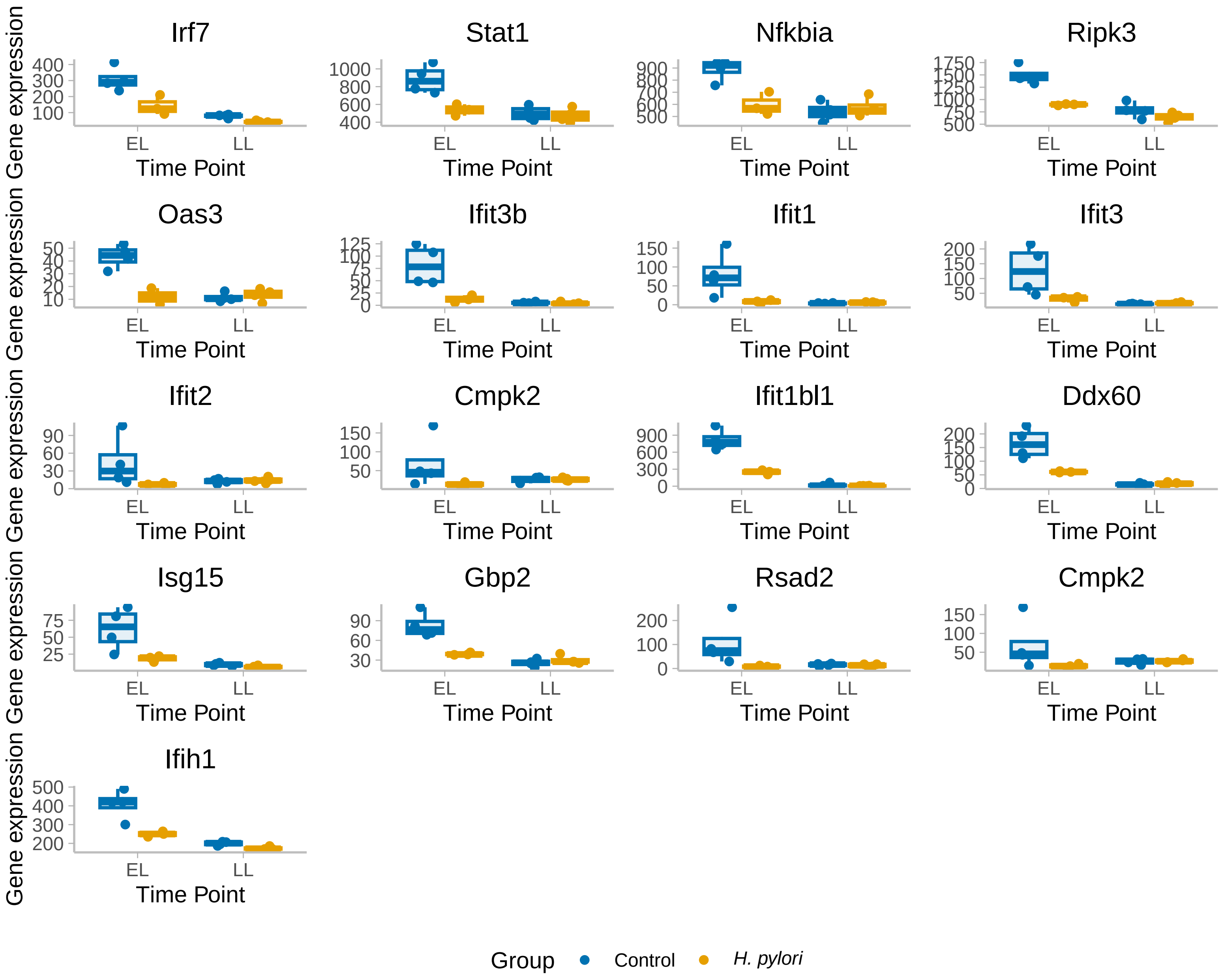

### Fig.s7

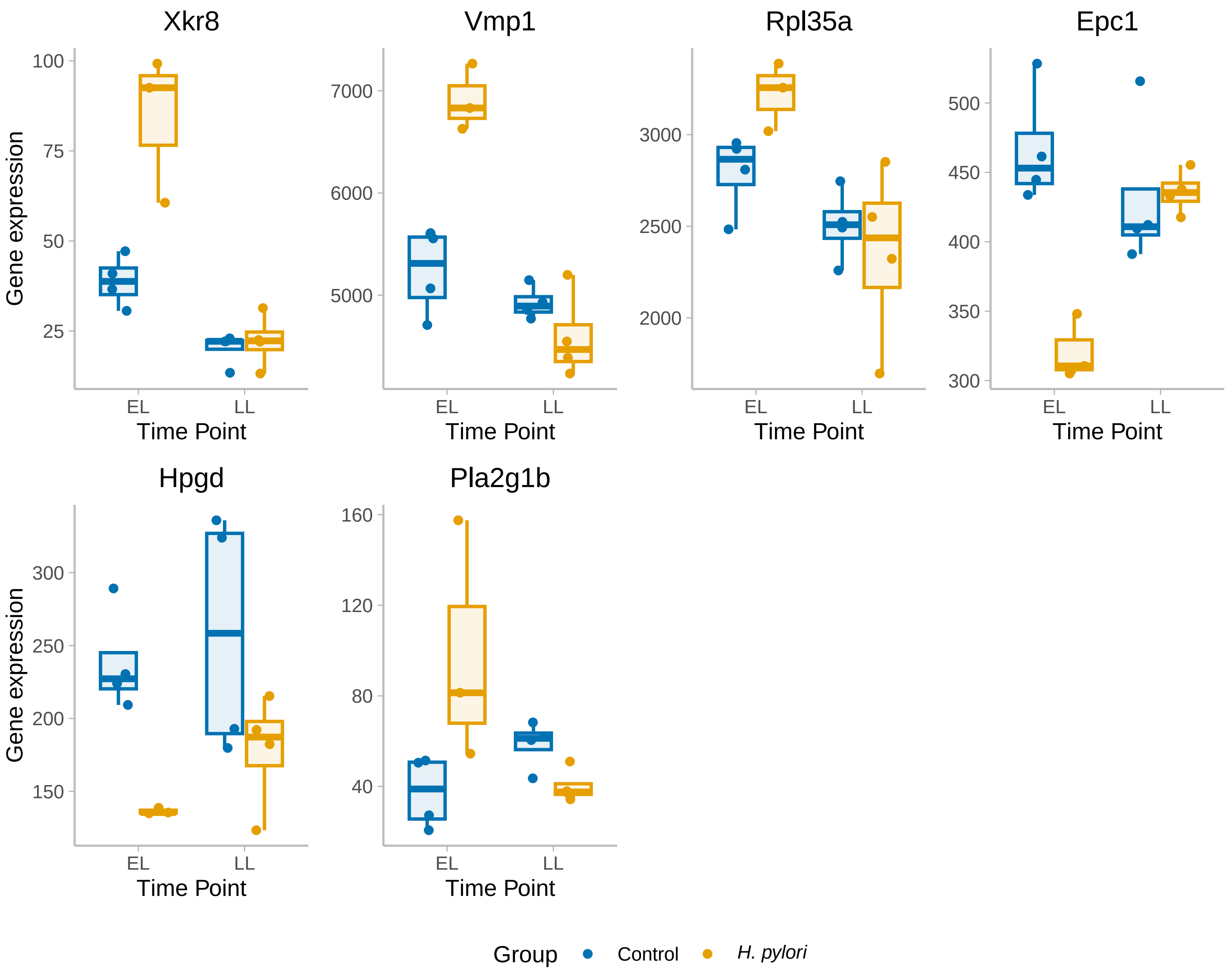

### Fig.s8

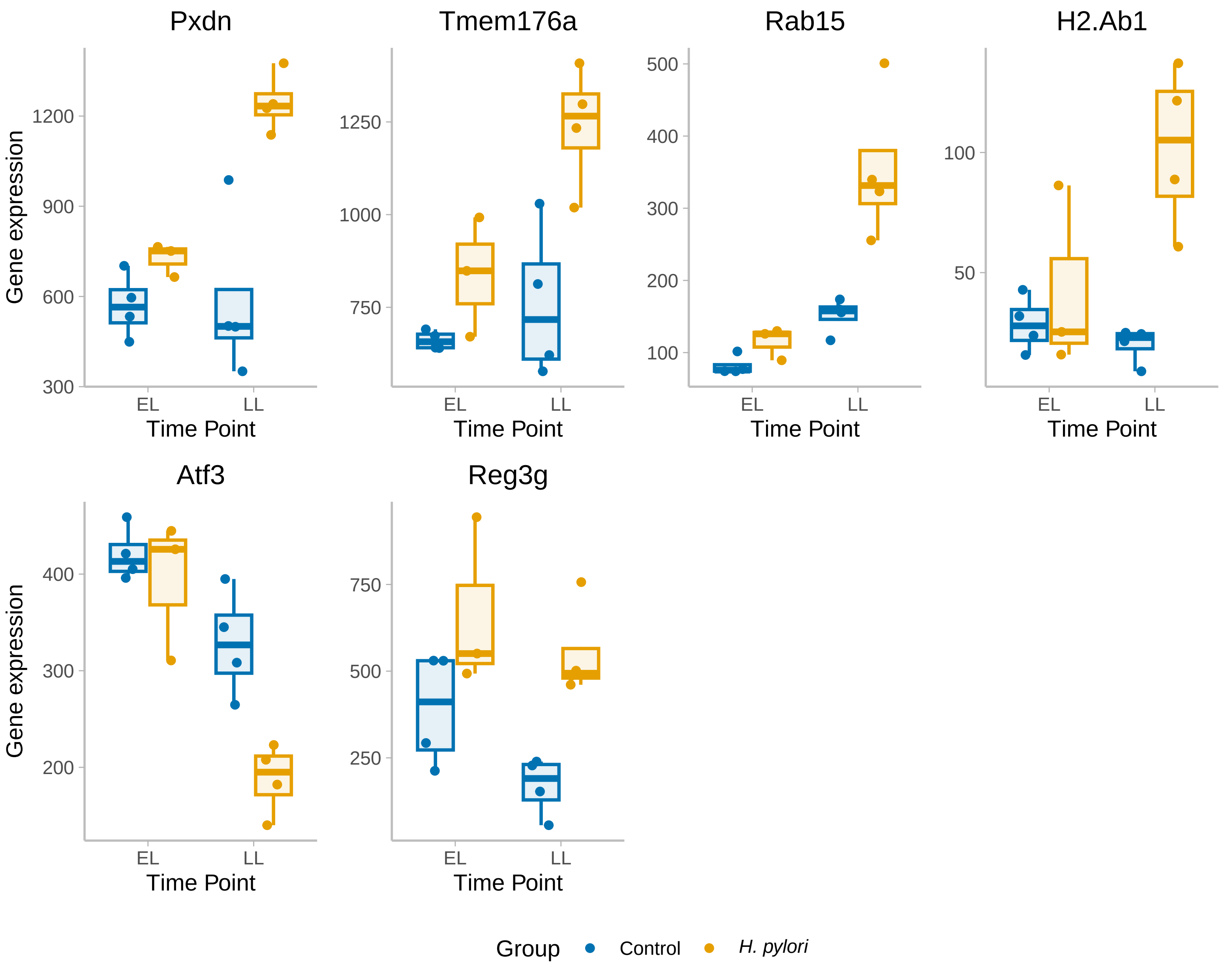
